# ATF4 regulates neuronal death in models of Parkinson’s Disease

**DOI:** 10.1101/783795

**Authors:** Matthew D. Demmings, Gillian N. Petroff, Heather E. Tarnowski-Garner, Sean P. Cregan

## Abstract

Parkinson’s Disease (PD) is characterized by the loss of dopaminergic neurons in the substantia nigra resulting in severe motor impairments. However, the mechanisms underlying this neuronal loss remain largely unknown. Activating Transcription Factor 4 (ATF4), a key mediator of the Integrated Stress Response (ISR), is a transcription factor that during prolonged activation can induce the expression of pro-apoptotic target genes. Oxidative stress and ER stress have been implicated in PD and these factors are known to activate the ISR. In this study, we have determined, that both PD neurotoxins (MPP+ and 6-OHDA) and α-synuclein aggregation induced by pre-formed human alpha-synuclein fibrils (PFFs) cause sustained upregulation of ATF4 expression in mouse primary cortical and mesencephalic neurons. Furthermore, we demonstrate that PD neurotoxins induce the expression of the pro-apoptotic factors Chop, Trb3 and Puma in an ATF4-dependent manner. Importantly, using neurons derived from ATF4 +/+ and ATF4 -/- mice, we demonstrate that ATF4 promotes neuronal apoptosis and dopaminergic cell loss in cellular models of PD. Finally, we demonstrate that the eIF2α kinase inhibitor C16 suppresses MPP+ and 6-OHDA induced ATF4 activation and protects against PD neurotoxin induced neuronal death. Taken together these results indicate that ATF4 is a key regulator of dopaminergic cell death induced by PD neurotoxins and pathogenic α-synuclein aggregates and highlight the ISR as a potential therapeutic target in PD.

## Introduction

Parkinson’s disease (PD) is the second most common neurodegenerative disorder and is characterized by distinct motor symptoms including bradykinesia, postural rigidity and resting tremor (Postuma et al., 2015). The motor impairments in PD result from the progressive loss of dopaminergic neurons in the substantia nigra pars compacta and the associated depletion of striatal dopamine (Damier, Hirsch, Agid, & Graybiel, 1999). A hallmark of PD is the appearance of intracellular protein inclusions called Lewy Bodies in which aggregates of misfolded α-synuclein are a major component (Spillantini et al., 1997). Although the majority of PD cases are sporadic, risk genes including PARK1 (α-synuclein), PARKIN, PINK1, DJ-1 and LRRK2, have been identified in familial forms of the disease (reviewed by Zeng, Geng, Jia, Chen, & Zhang, 2018). Unfortunately, there is no cure for PD and while current treatments temporarily ameliorate some of the clinical symptoms, they do not mitigate the underlying degenerative processes.

Numerous studies implicate oxidative stress and ER stress as major factors contributing to dopaminergic neuronal cell death in PD. Indeed, markers of oxidative damage including DNA damage, lipid peroxidation and aggregates of oxidized proteins have been reported in post-mortem brain tissue of PD patients (Baba et al., 1998; Bosco et al., 2006). The role of oxidative stress is also supported by the finding that environmental toxins (paraquat, rotenone) and neurotoxic agents (MPTP, 6-OHDA) can inhibit mitochondrial function and increase the production of reactive oxygen species cause loss of dopaminergic neurons in rodents, non-human primates and humans (Bove, Prou, Perier, & Przedborski, 2005). Furthermore, several familial-linked PD genes including Parkin, Pink1 and DJ-1 have been implicated in the regulation of mitochondrial function, mitophagy and ROS production (Clark et al., 2006; Geisler et al., 2010; Lee, Nagano, Taylor, Lim, & Yao, 2010; Narendra et al., 2010). In addition to oxidative stress, deficits in cellular proteostasis pathways associated with the accumulation and aggregation of misfolded proteins such as α-synuclein is thought to contribute to neurodegeneration in PD (Kaushik & Cuervo, 2015). One component of the proteostasis network that is affected in PD is the folding capacity of the endoplasmic reticulum (ER) resulting in ER stress and activation of the unfolded protein response (UPR) (Mercado, Castillo, Soto, & Sidhu, 2016; Michel, Hirsch, & Hunot, 2016). Indeed, markers of ER stress are commonly observed in post-mortem brain tissue derived from PD patients as well as in a variety of cellular and animal models of PD (Colla et al., 2012; Heman-Ackah et al., 2017; Holtz & O’Malley, 2003; Hoozemans et al., 2007; Mercado et al., 2018b; Ryu et al., 2002; Slodzinski et al., 2009).

The Integrated Stress Response (ISR) is a cell signaling pathway that is activated in response to diverse stress stimuli including oxidative stress and ER stress (Pakos-Zebrucka et al., 2016). Four kinases have been implicated in the initiation of the ISR including PERK (PKR-like ER kinase), PKR (protein kinase double stranded RNA-dependent), GCN2 (general control non-depressible-2), and HRI (heme-regulated inhibitor) (Donnelly, Gorman, Gupta, & Samali, 2013). The ISR kinases are activated in a stress specific manner and phosphorylate the translation initiation factor eIF2α (Lu et al., 2004). Phosphorylation of eIF2α results in attenuation of general protein translation but permits the selective translation of stress-induced mRNAs including that of Activating Transcription Factor 4 (ATF4) a member of the ATF/CREB family of basic-region leucine zipper transcription factors (Singleton & Harris, 2012). ATF4 regulates the expression of genes involved in amino acid metabolism and redox homeostasis that promote cell recovery (Harding et al., 2003). However, upon sustained activation ATF4 can also induce the expression of factors that have been implicated in promoting cell death including CHOP and TRB3 (Fawcett, Martindale, Guyton, Hai, & Holbrook, 1999; Ohoka, Yoshii, Hattori, Onozaki, & Hayashi, 2005; Zinszner et al., 1998). Induction of ATF4 has been reported in cellular and animal models of PD as well as in post-mortem brain tissue of PD patients (Mercado et al., 2018a; Ryu et al., 2002; Sun et al., 2013). However, the role of ATF4 in regulating neuronal survival remains controversial. Therefore, in the present study we investigated the role of ATF4 in primary neurons exposed to PD neurotoxins (MPP+ and 6-OHDA) and α-synuclein preformed fibrils.

## Materials and Methods

### Animals

All animal procedures were performed as per guidelines set by the Animal Care Committee at Western University, in accordance with the Canadian Council on Animal Care. Mice carrying an ATF4-null mutation were obtained from Drs. Tim Townes (University of Alabama, Birmingham, AL). All mice were maintained on a C57BL/6 background. Animals were genotyped as previously described (Masuoka & Townes, 2002).

### Primary cortical and mesencephalic neuron culture

Cortical neurons were dissociated from embryonic days 14.5-15.5 embryo’s and cultured in Neurobasal plus media (ThermoFisher, #A35829-01) supplemented with B27 plus (ThermoFisher, #A35828-01), 0.5x Glutamax (ThermoFisher, #35050-061), and 50U/mL penicillin: 50 μg/mL streptomycin (PenStrep; ThermoFisher, #15140-122). Mesencephalic neurons were obtained using a protocol modified from Gaven, Marin, and Claeysen, (2014) and cultured in the same media conditions as cortical neurons.

### Drugs

Drug treatments were initiated when neurons were 5-7 DIV. 1-methyl-4-phenylpyridinium iodide (MPP+; Millipore Sigma #D048), 6-Hydroxydopamine hydrobromide (6-OHDA; Millipore Sigma #H116), Thapsigargin (TG; Millipore Sigma #T9033), 6,8-Dihydro-8-(1*H*-imidazol-5-ylmethylene)-7*H*-pyrrolo[2,3-*g*]benzothiazol-7-one [C16] (Tocris #5382) were prepared in DMSO or ddH20 and diluted in neuron culture media immediately before administration to cultures at indicated concentrations.

### α-synuclein preformed fibrils

Human α-synuclein monomers (Proteos #RP-003) were used to generate fibrils based on protocol from Volpicelli-Daley, Luk, & Lee (2014). Briefly, monomers were shaken at 1000 rpm (37°C) for 7 days to obtained preformed fibrils, aliquoted and stored at −80°C. Aliquots were then thawed at room temperature immediately before use. PFFs were diluted in warm medium and sonicated with a probe tip at 10% power for 30 seconds (0.5 seconds on, 0.5 seconds off). Sonicated fibrils were added to neuronal culture at a final concentration of 5 µg/mL.

### Cell death and survival assays

Neuronal apoptosis was measured by visualizing nuclear morphology in Hoechst 33342 stained cells as described previously (Cregan et al., 2002). Briefly, neurons were fixed using Lana’s Fixative (4% paraformaldehyde, 0.2% picric acid) for 30 minutes, washed in PBS and stained with Hoechst 33342 (Invitrogen #H1399) at a concentration of 0.5 µg/mL. The fraction of cells exhibiting an apoptotic nuclear morphology characterized by pyknotic and/or fragmented nuclei containing condensed chromatin was scored by an individual blinded to the treatments. Neuronal survival was assessed via Calcein-AM/ Ethidium-Homodimer (Live/Dead) staining (Invitrogen #L3224). Dye was diluted in warmed neuron medium immediately before use. Calcein-AM (1µM) and Ethidium Homodimer (3µM) was added for 10 minutes to determine the ratio of live (Calcein-AM positive) to dead (Ethidium Homodimer positive) cells. In both assays, neurons were visualized by fluorescent microscopy (Olympus) and images were captured using a CCD camera (Q-imaging) and Northern Eclipse software (Empix Imaging).

### Real-time quantitative PCR (RTqPCR) and quantitative PCR (qPCR)

RNA was isolated via Trizol as per guidelines set by manufacturer (Invitrogen # 15596018) and 40ng was used in one-step Sybr green reverse transcription (RT)-PCR protocol (Qiagen #204154). RT-PCR was carried out in a CFX Connect Real Time System (Biorad) and changes in gene expression were determined by the Δ(ΔCt) method using S12 transcript for normalization. Values reported for RTqPCR are fold increase in mRNA levels in treated samples relative to paired untreated controls for transcript of interest. RT-PCR amplifications were carried out as follows: 50°C for 10 min, 95°C for 5 min, 95°C for 10 sec, and 60°C for 30 sec. Primer sequences used for amplification available upon request.

### Western blot analysis

To obtain whole-cell lysates, neurons were incubated in lysis buffer (RIPA buffer (Millipore Sigma, #R0278) supplemented with phosphatase inhibitor cocktail (Millipore Sigma, #P5726) and protease inhibitor cocktail (Millipore Sigma, #P8340)) on ice for 30 minutes and soluble extract was recovered via centrifugation. Protein concentrations were determined by BCA assay (ThermoScientific, #23225). 40µg of protein was separated on 12% SDS-PAGE and transferred to PVDF membrane. Membranes were then blocked for 1hr at room temperature in 5% Milk prepared in TBST (10 mM Tris, 150 mM NaCl, 0.1% Tween 20). Membranes were the incubated overnight with primary antibodies against ATF4 (1: 5000; Abcam #184909), Cyclophilin B (1: 5000; Abcam #178397), and Cleaved Caspase 3 (1: 1000; Cell Signaling Technologies #9661) at 4°C. Membranes were then washed in TBST and incubated at room temperature with HRP-conjugated goat anti-rabbit secondary (1: 10000; BioRad #1706515) for 1 hour. Membranes were then washed again and developed via enhanced chemiluminescence (BioRad #1705061).

### Immunocytochemistry

Briefly, neurons were washed with ice cold PBS containing Na_3_VO_4_ (1mM; Millipore Sigma #S-6508)) and NaF (25mM; Millipore Sigma # S-6776). Neurons were then fixed in Lana’s fixative (4% paraformaldehyde, 0.2% picric acid) and washed in PBS. Cells were then permeabilized in ice cold 100% MeOH for 10 mins and washed in PBS. Neurons were then blocked in 2% BSA for 1hour and incubated with primary antibodies against MAP2 (1: 500; Abcam #11267), ATF4 (1:500; Santa Cruz #SC-200), Cleaved Caspase 3 (1: 400; Cell Signaling Technologies #9661), Tyrosine Hydroxylase (1:500; Millipore Sigma #AB152), and p-serine 129 α-synuclein (1:500; Abcam 59264) overnight at 4°C. The next day, cells were washed with PBS and incubated with appropriate AlexFluor (Invitrogen) for 2 hours at room temperature. Neurons were then washed in PBS and Hoechst stained. Coverslips were mounted onto glass microscope slides (VWR #48311-600) using ProLong Gold mounting glue (Invitrogen #P36930). Images were obtained using previously described fluorescent microscopy, EVOS Imaging System (ThermoFisher), and confocal images were taken on SP8 microscope (Lieca Microsystems, Germany).

### Mitochondrial Superoxide (MitoSox) Staining

5mM of Mitosox dye (Molecular Probes #M36008) was prepared as per manufacturer’s instructions. Dye was then diluted in warmed neuron medium and added to cultures at final concentration of 200nm for 2 hours at 37°C / 5% C0_2_. Neurons were then washed with PBS and fixed with Lana’s fixative. Subsequently, neurons were incubated with antibodies against MAP2 and appropriate secondary, then mounted and imaged as described above.

### Statistical Analysis

Data are reported as mean ± standard error of the mean (SEM). Statistical analyses were performed using GraphPad Prism version 8.2 for Mac (GraphPad Software, USA). Statistical significance was considered was set a p <0.05 and determined via t-test or ordinary one-way and two-way ANOVA with Tukey and Sidak post-hoc tests. The ‘n’ value indicates the number of independent neuron culture experiments or the number of embryos of each genotype from which neurons were dissociated from.

## Results

### PD neurotoxins MPP+ and 6-OHDA cause sustained activation of ATF4 in neurons

The neurotoxins 6-hydroxydopamine (6-OHDA) and 1-methyl-4-phenyl-pyridium (MPP+) cause death of a dopaminergic neurons *in vitro* and *in vivo* and have been widely used to investigate cellular mechanisms involved in dopaminergic neuron degeneration in PD (Przedborski & Ischiropoulos, 2005). These neurotoxins are known to induce mitochondrial dysfunction and to produce ROS that lead to neuronal injury and cell death (Fallon, Matthews, Hyman, & Beal, 1997; Lotharius, Dugan, & O’Malley, 1999). Therefore, to investigate the role of ATF4 in PD we initially examined whether these PD-mimetics induce ATF4 expression in primary cortical neurons. As shown in Figures 1 A & 1B, both MPP+ and 6-OHDA treatments resulted in a robust (>5-fold) increase in ATF4 expression that persisted for at least 16-24 hours.

**Figure 1.**
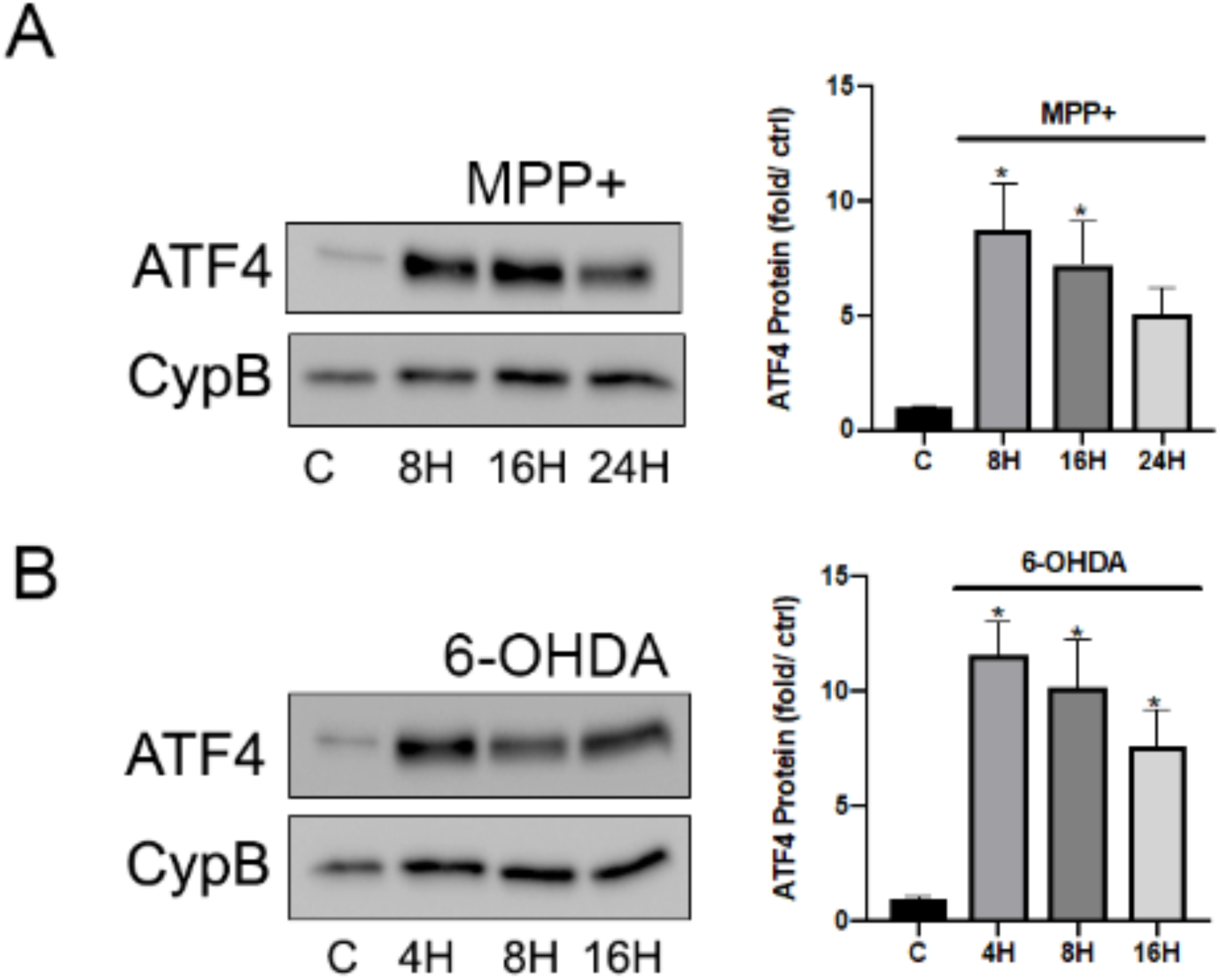
PD neurotoxins MPP+ and 6-OHDA cause sustained induction of ATF4 in neurons. Cortical neurons were treated with MPP+ [50 µM] (A) or 6-OHDA [10 µM] (B) and at the indicated timepoints ATF4 protein levels were assessed by Western blot analysis and quantified by densitometry (n=3; *p<0.05). C, Cortical neurons were treated with MPP+ [50 µM] or 6-OHDA [10 µM] for 8h and mRNA levels of ATF4 was determined by quantitative RT-PCR. Induction was normalized to S12 mRNA levels and is reported as fold increase over untreated neurons (n=3; p>0.05)

### ATF4 is required for transcriptional induction of the pro-apoptotic genes CHOP, Trib3, and PUMA during exposure to PD neurotoxins

ATF4 is a stress responsive transcription factor that can induce the expression of genes involved in amino acid metabolism and redox homeostasis. However, during prolonged stress ATF4 has also been implicated in the regulation of pro-apoptotic genes (Harding 2003; Zinzsner 1998). CHOP, Trib3, and PUMA are known to have pro-apoptotic activity and have been previously reported to be induced by PD neurotoxins and to contribute to subsequent neuronal death (Aimé et al., 2015; Bernstein, Garrison, Zambetti, & O’Malley, 2011; Bernstein & O’Malley, 2013; Silva et al., 2005). Therefore, we sought to determine whether the induction of these pro-death factors by PD neurotoxins is regulated by ATF4. To address this, we examined mRNA levels of Chop, Trib3 and Puma in ATF4 +/+ and ATF4 -/- cortical neurons following treatment with PD toxins. Treatment of ATF4+/+ neurons with MPP+ or 6-OHDA resulted in a marked increase in the expression of CHOP, Trib3 and Puma mRNAs (Figure 2A and 2B). However, the increase in expression of these pro-apoptotic factors in response to PD neurotoxins was largely attenuated in ATF4-/- neurons. Taken together these results indicate that sustained activation of ATF4 is required for transcriptional induction of key pro-apoptotic target genes by PD neurotoxins.

**Figure 2.**
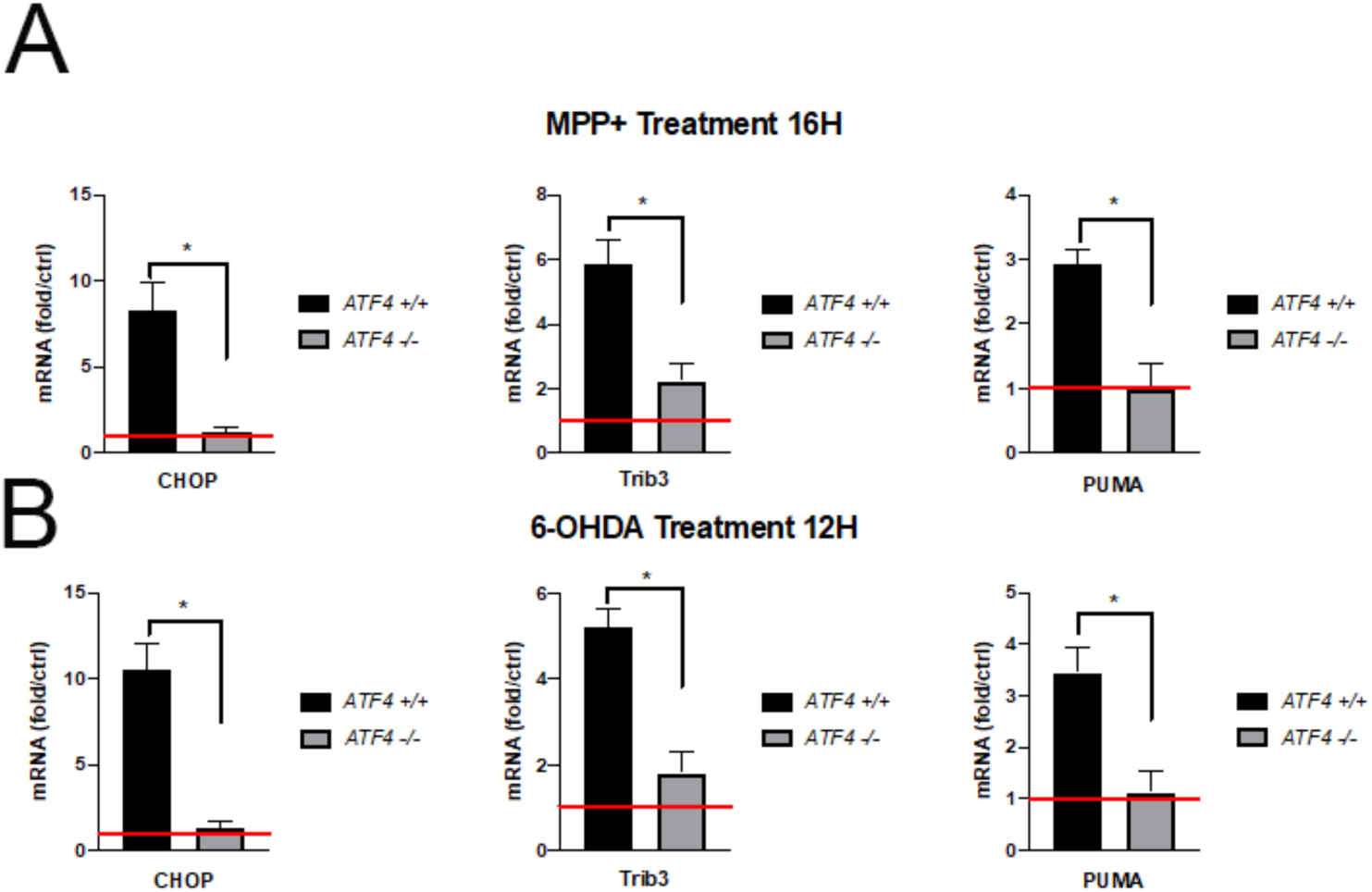
PD mimetics trigger ATF4-dependent transcription of the pro-apoptotic genes Chop, Trb3 and Puma. Cortical neurons derived from ATF4-wildtype and ATF4-null littermates were treated with (A) MPP+ [50 µM] for 16h or (B) 6-OHDA [10 µM] for 12h and mRNA levels of indicated transcripts were determined by quantitative RT-PCR. Induction was normalized to S12 mRNA levels and is reported as fold increase over untreated neurons (n=4; *p<0.05).

### ATF4 is required for PD-neurotoxin induced neuronal apoptosis

Having established that ATF4 promotes the transcriptional induction of several pro-apoptotic target genes following exposure to PD neurotoxins we next investigated whether ATF4 is required for MPP+ and 6-OHDA induced neuronal death. However, since a previous study had indicated that ATF4 can regulate the expression of genes involved in redox homeostasis we wanted to first determine whether the level of oxidative stress induced by PD neurotoxins is affected in ATF4-deficient neurons. To address this, we treated ATF4+/+ and ATF4-/- neurons with MPP+ or 6-OHDA for 8 hours and then assessed the level of mitochondrial superoxide production using MitoSOX Red assay. The MitoSOX probe targets the mitochondria and fluoresces red upon oxidation by superoxide free radicals. As shown in figure 3, both MPP+ and 6-OHDA treatments produced significant increases in superoxide levels as determined by the increased MitoSOX Red fluorescence intensity. Furthermore, the level PD-neurotoxin induced superoxide formation was not significantly different between ATF4+/+ and ATF4-/- neurons (Figure 3A and 3B). It was also noted that wildtype and ATF4-deficient neuronal cultures exhibited a similar, healthy neuronal morphology as depicted by MAP2 immunostaining (Figure 3B). Next, to determine whether ATF4 is required for PD-neurotoxin induced neuronal apoptosis we treated ATF4+/+ and ATF4-/- neurons with MPP+ or 6-OHDA and then quantified the level of apoptosis as a function of time by assessing nuclear morphology following Hoechst staining. As depicted in Figure 4A, treatment of wild-type neurons with MPP+ or 6-OHDA over a 48-hour period resulted in a substantial increase in the fraction of cells exhibiting an apoptotic morphology characterized by chromatin condensation and/or nuclear fragmentation. In contrast, in ATF4-deficient neurons MPP+ and 6-OHDA treatment resulted in only a modest increase in apoptosis over this same time frame, and the level of apoptosis was found to be significantly lower than in ATF4+/+ neuronal cultures at both 24 hours and 48 hours post-treatment (Figure 4A). Consistent with the apoptotic counts we also found that the level of cleaved (active) Caspase-3 induced by MPP+ and 6-OHDA treatments was markedly reduced in ATF4-/- neurons as compared to ATF4+/+ neurons (Figure 4B and 4C). Next to determine whether ATF4-deficient neurons remained viable following treatment with PD-neurotoxins and were not just dying by a non-apoptotic mode of cell death we assessed neuronal survival in ATF4 +/+ and ATF4 -/- using Calcein-AM/ethidium homodimer staining (Live/Dead assay). As shown in Figure 4D, MPP+ and 6-OHDA treatment markedly reduced cell viability in ATF4+/+ neuronal, but not in ATF4-/- neuronal cultures indicating that ATF4-deficient neurons remain viable.

**Figure 3.**
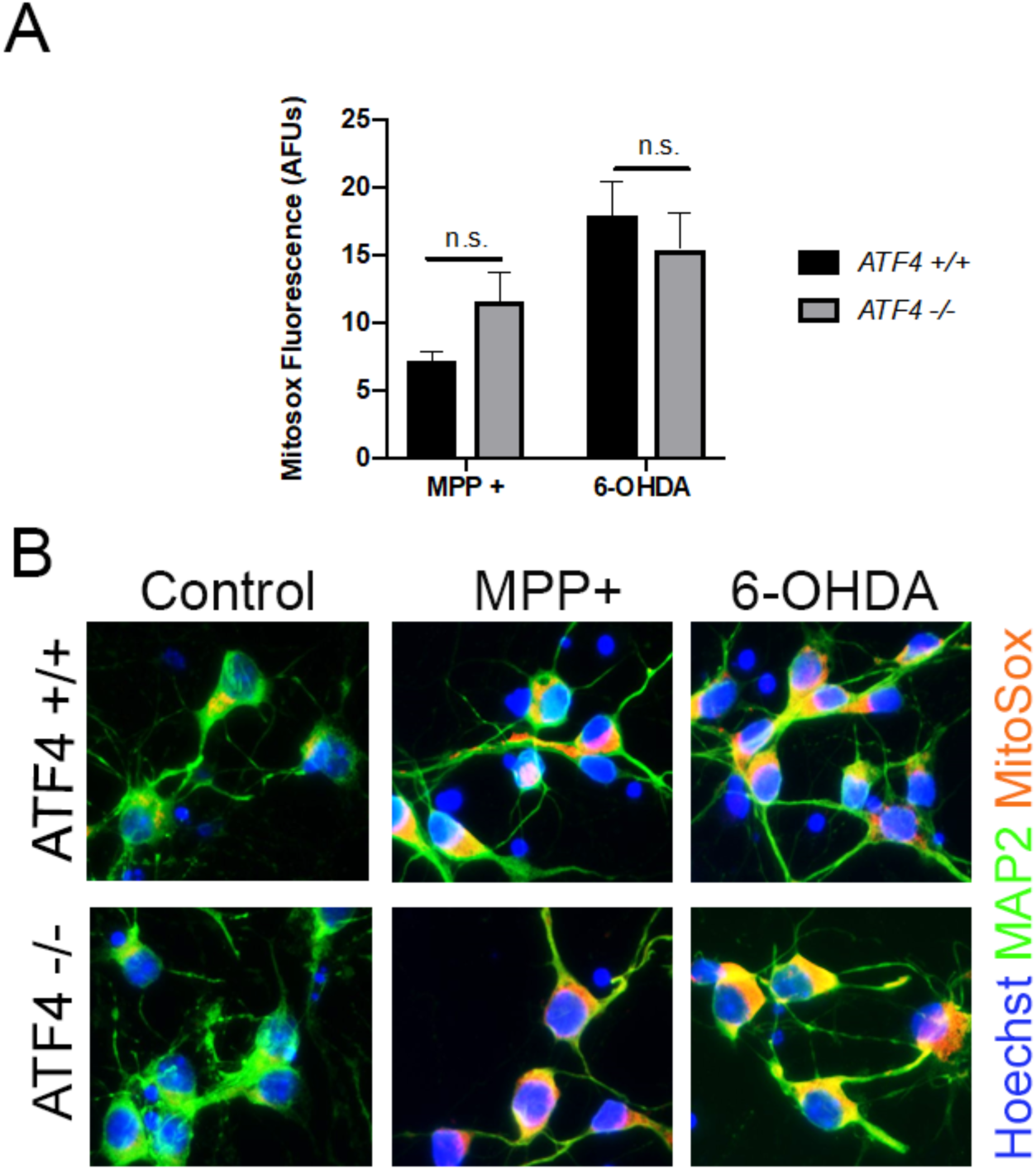
Oxidative stress produced by MPP+ and 6-OHDA is not dependent on ATF4 expression. B, Cortical neurons were treated with MPP+ [50 µM] or 6-OHDA [10 µM] and mitochondrial superoxide accumulation was measured and quantified at 8h (n=3; p>0.05). B, Representative images of MAP2-positve (green) neuronal cultures and accumulated superoxide as indicative by MitoSox (red) fluorescence.

**Figure 4.**
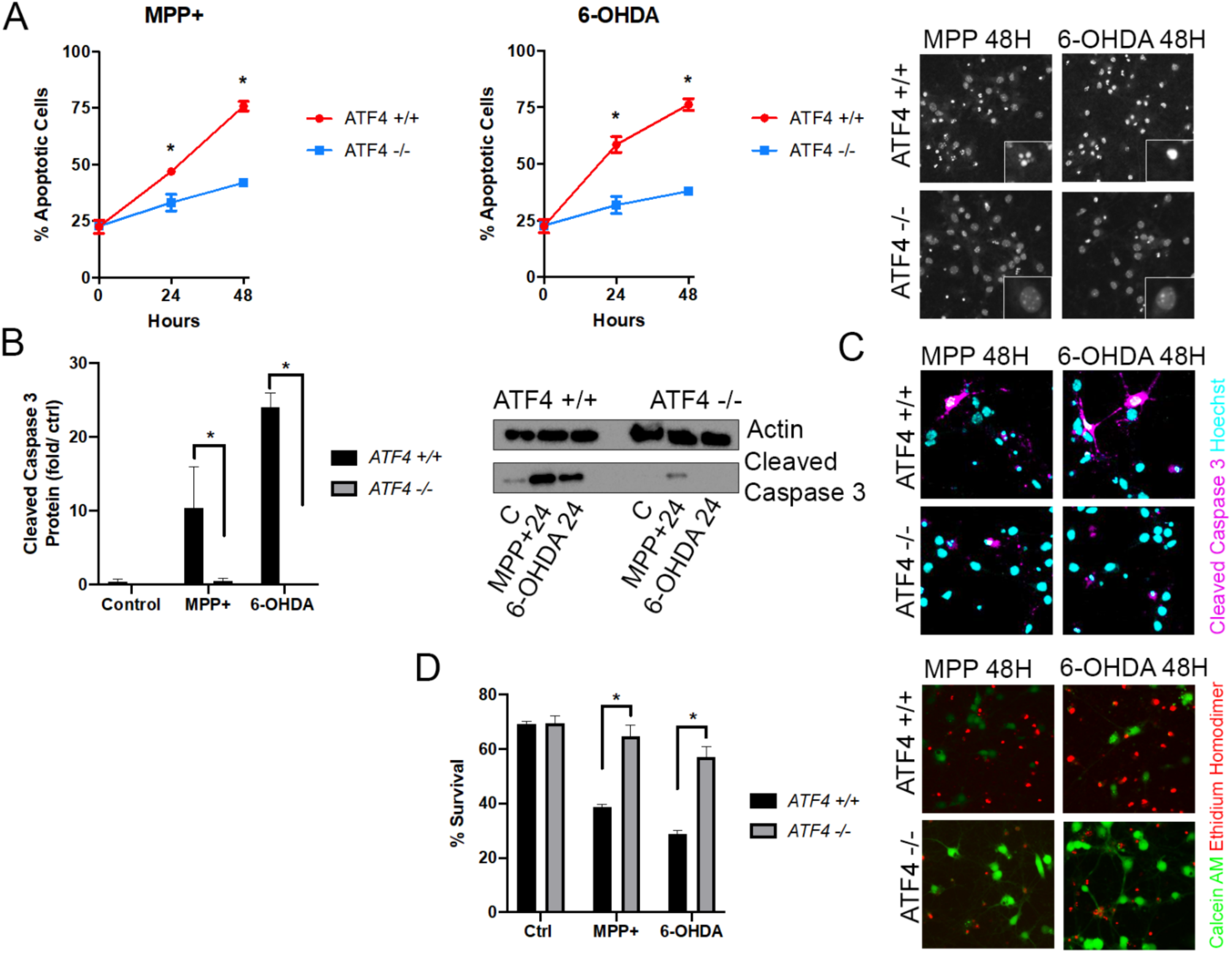
ATF4 is required for PD toxin-induced neuronal apoptosis. A, cortical neurons derived from ATF4-wildtype and ATF4-null littermates were treated with MPP+ [50 µM] or 6-OHDA [10 µM] and the percentage of apoptotic cells was determined by Hoechst 33442 staining at 24 and 48h. (n ≥ 5; *p<0.05). Representative images of Hoechst 33442 stained neurons captured at 48h. B, cortical neurons derived from ATF4 wild-type and ATF4-null littermates were treated with MPP+ [50 µM] or 6-OHDA [10 µM] and levels of cleaved caspase 3 were assessed at 24H by Western blot analysis and quantified by densitometry (n=3; *p<0.05). C, Representative immunofluorescent images of cleaved caspase 3 in ATF4-wildtype and ATF4-null neuronal cultures treated with MPP+ (50 µM) or 6-OHDA (10 µM) for 24h. D, cortical neurons derived from ATF4-wildtype and ATF4-null littermates were treated with MPP+ [50 µM] and 6-OHDA [10 µM] and survival was assessed by Calcein-AM/ Ethidium-Homodimer (live/dead) staining (n ≥ 4; *p<0.05). Representative fluorescent images of MPP+ and 6-OHDA treated ATF4-wildtype and ATF4-deficient neurons at 48H using live (green)/dead (red) assay.

Dopaminergic neurons of the substantia nigra are the primary neuronal population of neurons affected in PD. Therefore, we sought to determine whether ATF4 promotes cell death specifically in dopaminergic neurons exposed to PD toxins. To address this question, we generated mesencephalic cultures from the ventral midbrain region of ATF4+/+ and ATF4-/- mouse embryos and subsequently exposed them to MPP+ or 6-OHDA. Only a small portion of the neurons in these mesencephalic cultures are dopaminergic (DA) therefore In order to quantify the extent of DA neuron loss in these models, we quantified the number of neurons with positive immunoreactivity for tyrosine hydroxylase (TH), an enzyme responsible in the conversion of L-tyrosine to L-dihydroxyphenylalanine (L-DOPA) (Nagatsu, Levitt, & Udenfriend, 1964). As shown in Figure 5, MPP+ treatment resulted in the loss of 63.83% of the DA neurons in ATF4+/+ cultures as compared to only a 37.05% reduction in ATF4-/- cultures. Similarly, 6-OHDA treatment resulted in the loss of 52.39% of the DA neurons in ATF4+/+ cultures as compared to only a 12.95% reduction in ATF4-/- cultures. Taken together, these results indicate that ATF4 is necessary for PD neurotoxin induced cell death in dopaminergic neurons.

**Figure 5.**
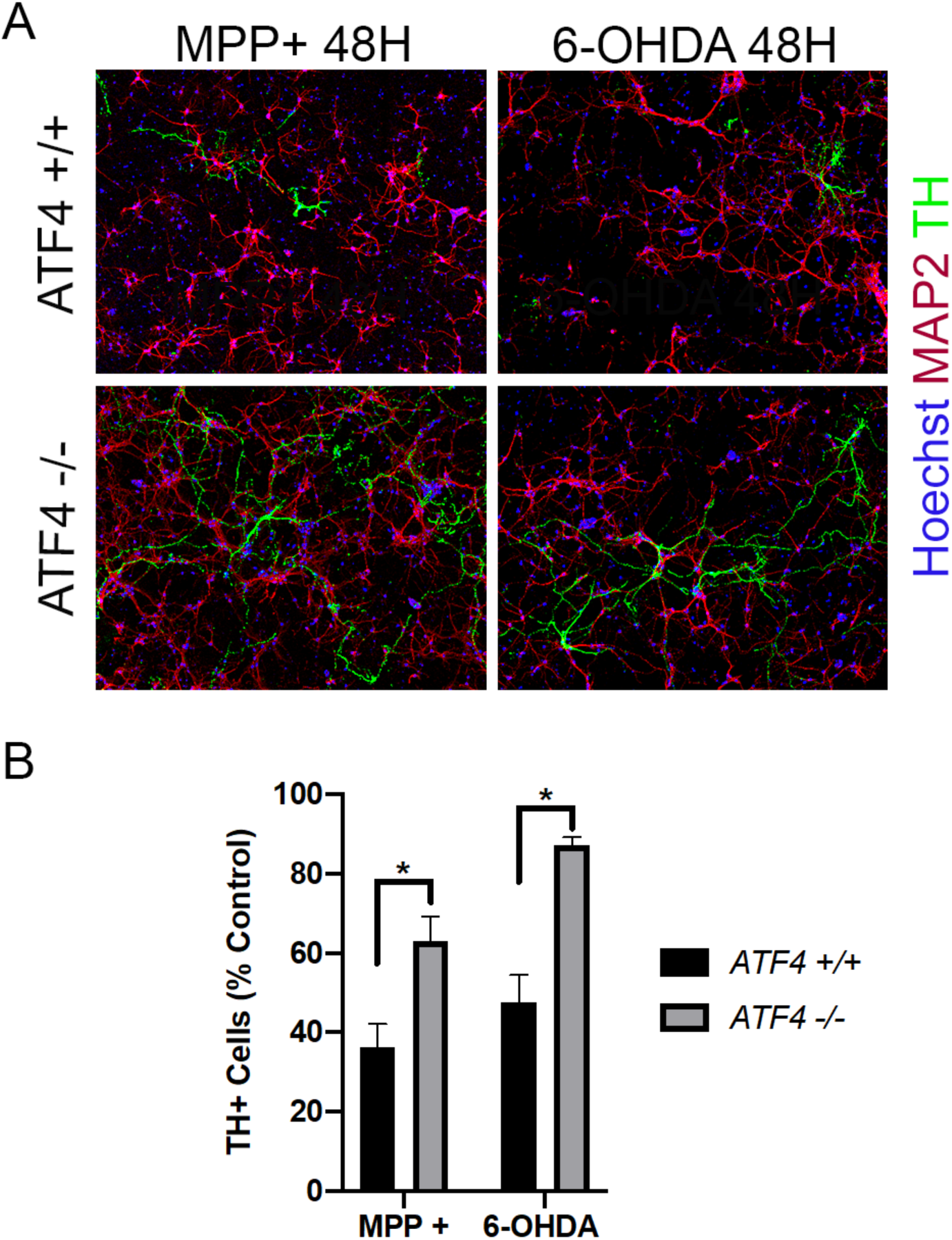
Dopaminergic neurons are conserved in ATF4-null mesencephalic cultures exposed to MPP+ or 6-OHDA. Mesencephalic neurons derived from ATF4-wildtype and ATF4-null littermates were treated with MPP+ [25 µM] or 6-OHDA [5 µM] for 48h and then immunostained for the dopaminergic neuron marker tyrosine hydroxylase (TH). A, Representative images showing increased numbers of residual dopaminergic neurons (TH+, green) in ATF4-/- mesencephalic cultures compared to ATF4+/+ cultures following treatment wih MPP+ or 6-OHDA. Cultures were co-stained for the general neuronal marker MAP2 and the nuclear stain Hoechst. B, Quantification of the total number of TH+ neurons in ATF4+/+ and ATF4-/- mesencephalic cultures treated with MPP+ (25 µM) or 6-OHDA (5 µM) for 48h. TH+ cell counts are reported as a percentage of untreated neurons from the same culture (n=3; *p<0.05).

### Human α-synuclein preformed fibrils induces ATF4 dependent neuronal loss

To complement our cellular neurotoxin models, we used human alpha-synuclein to elicit pathogenic aggregates in vitro to model the synucleinopathy present in PD. Previous studies have demonstrated that exogenous preformed **α**-synuclein fibrils (α-Syn PFFs) can be taken up into cultured neurons via endocytosis and induce aggregation of endogenous α-synuclein leading to the formation of intracellular inclusions that in turn, can lead to synaptic dysfunction and neuronal death (Volpicelli-Daley et al., 2011). Synuclein incorporated into pathogenic inclusions is highly phosphorylated at Ser129 (Fujiwara et al., 2002). Therefore, to validate this synucleinopathy paradigm we treated cortical neurons with α-Syn PFFs and confirmed the formation of phospho-Ser193-α-synuclein positive inclusions (Figure 6A). We then performed ATF4 immunostaining to determine whether α-Syn PFF induced lesions resulted in the induction of ATF4 expression. Indeed, as shown in Figure 6B, α-Syn PFF treated cultures exhibited a significant increase in nuclear ATF4 staining as compared to untreated neurons. We next sought to determined whether α-Syn PFF induced inclusions led to neuronal death, and if so whether this was dependent upon ATF4 expression. To address this question, we treated cortical neurons derived from ATF4+/+ and ATF4-/- mice with α-Syn PFFs and then assessed apoptosis by examining nuclear morphology. As shown in Figure 6C, α-Syn PFF treatment caused a significantly greater induction of apoptosis in ATF4+/+ neurons than in ATF4-/- neurons. Next, we probed mesencephalic cultures treated with preformed α-synuclein fibrils to examine whether loss of ATF4 affects pathogenic α-synuclein induced dopaminergic neuron death. A-Syn PFF induced synucleinopathy resulted in the loss of 81.65% of TH+ dopaminergic neurons in ATF4+/+ cultures, but only a 41.85% reduction in dopaminergic neurons in ATF4-/- cultures (Figure 6D). Taken together these results indicate that ATF4 is activated in response to pathogenic α-synuclein aggregation and promotes neuronal death.

**Figure 6.**
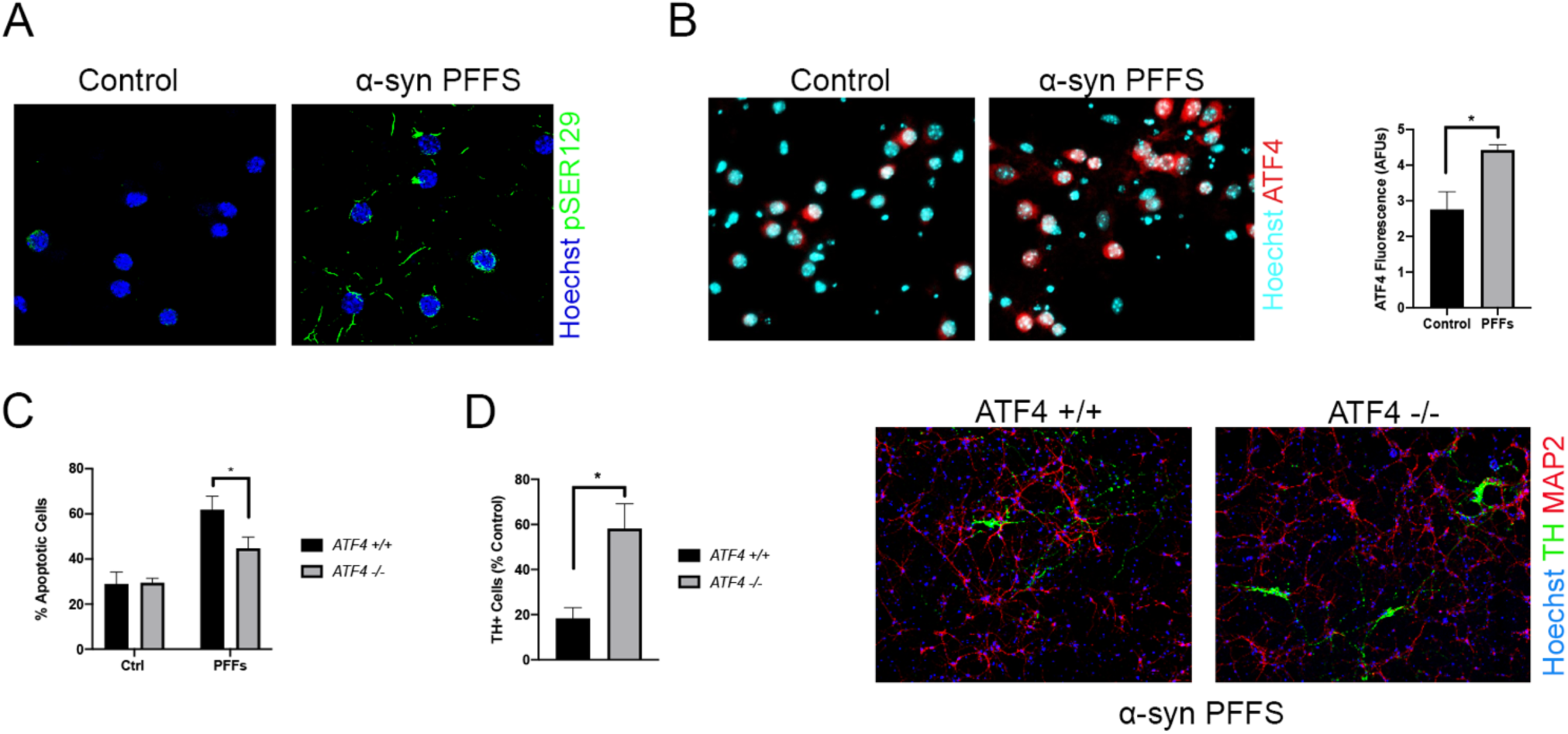
Human alpha-synuclein pre-formed fibrils (α-Syn PFFs) induce ATF4-dependent neuronal death. Cortical neurons were treated with α-Syn PFFs [5µg/ml] or left untreated (control) for 12 days. A, Representative images showing pathogenic intracellular aggregates of α-Synuclein identified by p-Serine129-α-synuclein immunoreactivity (green). B, Representative images of ATF4 immunostaining (red) and quantification of ATF4 immunofluorescence intensity (n=3; *p<0.05). C, Cortical neurons derived from ATF4-wildtype and ATF4-null littermates were treated with α-Syn PFFs [5 µg/ml] for 14 days and the percentage of apoptotic cells was determined by Hoechst staining (n=4; *p<0.05). D, mesencephalic neurons derived from ATF4-wildtype and ATF4-null littermates were treated with PFFs [5 µg/ml] for 14 days (n=3; *p<0.05). Representative images of dopaminergic (TH+) neurons (green) in wildtype and ATF4-deficient mesencephalic cultures exposed to α-Syn PFFs for 14 days.

### The eIF2α kinase inhibitor C16 inhibits ATF4 activation and protects against neuronal death in cellular models of PD

Oxidative stress and ER stress have been implicated in PD and are known to trigger the Integrated Stress Response and subsequent induction of ATF4 through the activation of eIF2α kinases. Given our findings that ATF4 is activated in PD models and promotes neuronal death we reasoned that pharmacological inhibition of eIF2α kinases that contribute to ATF4 activation in response to oxidative stress and ER stress may be neuroprotective in PD. In preliminary experiments we found that the imidazole-oxindole PKR inhibitor commonly known as C16 was able to inhibit ATF4 induction in neurons following treatment with the oxidative stressor arsenite as well as the ER stress inducing agent thapsigargin (data not shown). Therefore, we tested the effects of this eIF2α kinase inhibitor in cellular models of PD. We first investigated whether C16 could inhibit PD neurotoxin induced ATF4 activation. As shown in figure 7A, C16 administration significantly reduced the levels of ATF4 induced by both MPP+ and 6-OHDA treatments. We next investigated whether C16 could inhibit the induction of the ATF4-dependent pro-apoptotic target genes Chop and Trib3. Consistent with the ability of C16 to inhibit ATF4 induction, we found that the transcriptional induction of Chop and Trb3 mRNAs by MPP+ and 6-OHDA was significantly attenuated in the presence of C16 (Figure 7B). Given that C16 decreased ATF4 protein and represses transcriptional activation of its downstream targets, we next evaluated whether C16 is neuroprotective in PD neurotoxin paradigms. To test this, we treated cortical neurons with MPP+ or 6-OHDA in the presence or absence of C16 and then assessed the level of apoptotic cell death after 24 hours. As shown in Figure 7C, administration of C16 markedly reduced the level of apoptotic cell death induced by both MPP+ (63.24 vs 34.94%, p<0.05) and 6-OHDA (71.34% vs 45.81%, p<0.05).

**Figure 7.**
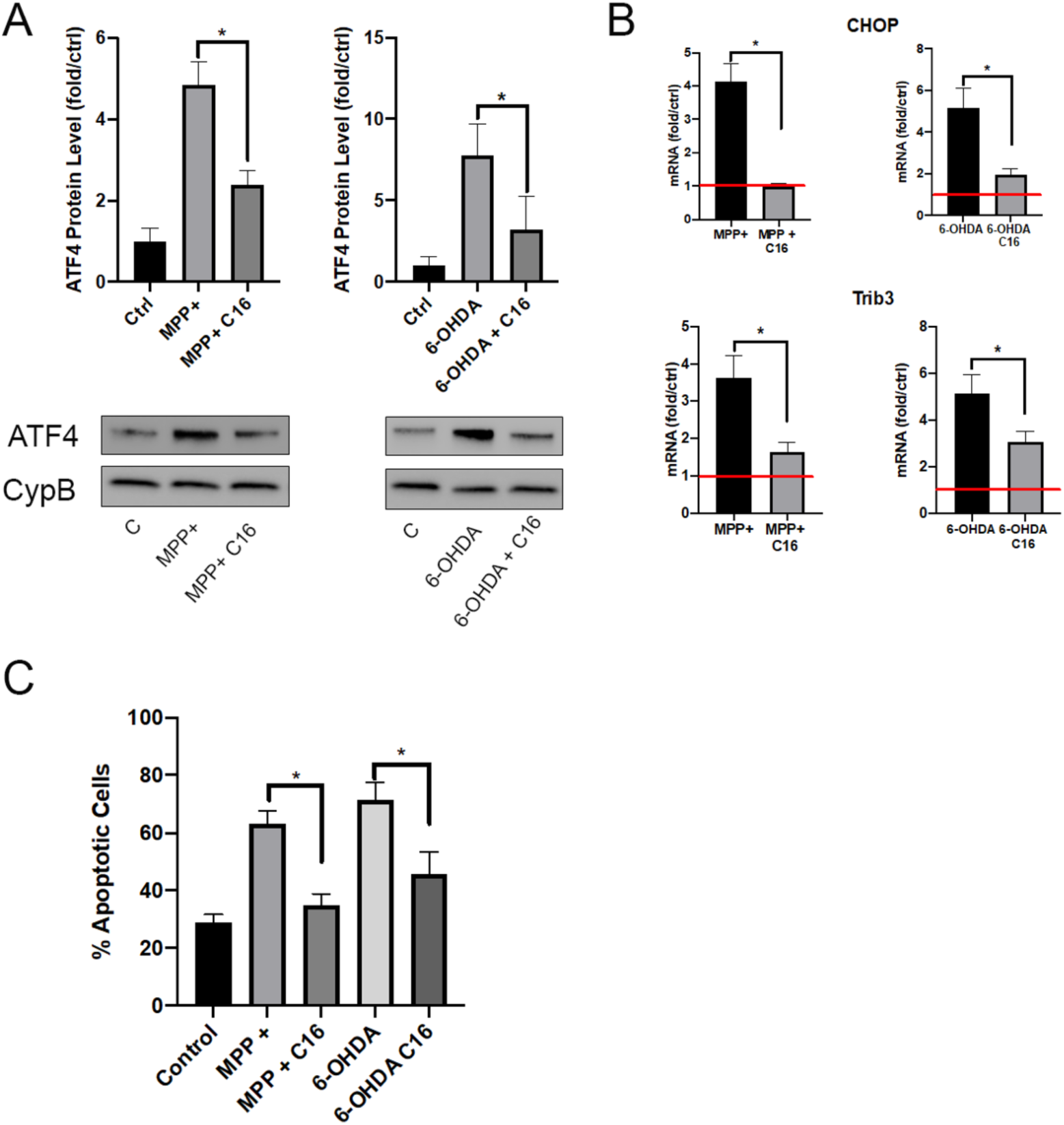
The eIF2a kinase inhibitor C16 attenuates PD neurotoxin induced ATF4 activation and neuronal cell death. A, cortical neurons were treated with MPP+ [50 µM] or 6-OHDA [10 µM] for 8h in the presence or absence of C16 [2 µM] and ATF4 protein levels were assessed by Western blot analysis and quantified by densitometry. ATF4 expression was normalized to Cyclophilin B levels (n=3; *p<0.05). B, mRNA levels of ATF4-dependent pro-apoptotic targets Chop and Trib3 were determined by quantitative RT-PCR 8h following treatment with MPP+ or 6-OHDA in the presence or absence of C16. Induction was normalized to S12 mRNA levels and is reported as fold increase over untreated neurons (n=5; *p<0.05). C, cortical neurons were treated with MPP+ [50 µM] or 6-OHDA [10 µM] in the presence or absence of C16 [2 µM] and the percentage of apoptotic cells was determined at 24 hours by Hoechst staining (n=3; *p<0.05). D, mRNA levels of UPR marker BiP in cortical neurons were determined by quantitative RT-PCR at 8H (n=3; *p<0.05) and spliced XBP1 patterns at 8h were assessed in triplicate via qPCR (representative image shown).

## Discussion

Parkinson’s disease is characterized by the loss of dopaminergic neurons, leading to a reduction of dopamine levels, which produce the cardinal motor symptoms of the disease. Currently, treatments are directed towards dopamine replacement therapies, which provide some symptomatic relief, however become less effective over time and do not attenuate the progression of the disease. Unfortunately, the underlying mechanism of neurodegeneration in Parkinson’s disease remains poorly understood. In this study, we have demonstrated that ATF4 induction in cellular models of PD triggers neuronal cell death through transcriptional activation of known pro-death genes and subsequent apoptotic processes. Specifically, we have determined that (a) ATF4 expression is increased following neuronal injury caused by PD neurotoxins (MPP+ and 6-OHDA) as well as by α-synuclein aggregates produced by preformed α-synuclein fibrils, that (b) ATF4 is transcriptionally active and promotes the expression of the pro-apoptotic target genes Trib3, CHOP, and PUMA, that (c) ATF4-deficient dopaminergic neurons are protected in cellular PD models, and importantly that (d) pharmacological inhibition of ATF4 attenuates neuronal loss in PD paradigms.

ATF4 is known to induce the expression of genes involved mitigating cellular stresses but during prolonged activation is also known to induce the expression of pro-apoptotic factors (Harding et al., 2003; Zinszner et al., 1998). Therefore, it is not surprising that ATF4 has been reported to have both pro-survival and pro-death effects in neurons. Specifically, ATF4 has been reported to promote resistance of HT22 and PC12 cells to glutamate induced oxytosis by inducing the expression of the cystine/glutamate antiporter-xCT (Lewerenz et al., 2011; Lewerenz & Maher, 2009). Conversely, in another study it was shown that ATF4-deficient cortical neurons are resistant to homocysteine induced oxidative stress and that overexpression of ATF4 is sufficient to promote neuronal death (Lange et al., 2008). Furthermore, we have previously demonstrated that ATF4-deficient neurons are resistant to ER stress but not DNA damage induced apoptosis (Galehdar et al., 2010). In this study we show that ATF4 functions as a pro-death effector in both neurotoxin and pathological α-synuclein models of PD. In agreement with our findings that ATF4 induction promotes dopaminergic neuronal loss, Gully et al., (2016) demonstrated that overexpression of ATF4 is sufficient to induce dopaminergic neuron loss in the substantia nigra *in vivo*. In contrast, it has been reported that in PC12 cells ATF4 expression induces Parkin expression and promotes cell survival in PD toxin paradigms (Sun et al., 2013). However, a more recent study by this group demonstrated that the ATF4 target Trib3 is induced in cellular models of PD and promotes neuronal death by decreasing Parkin levels in neurons (Aimé et al., 2015). Although our results demonstrate that ATF4 is a key regulator of neuronal death in cellular models of PD, future studies will be needed to evaluate the role of ATF4 in *in vivo* models of PD. However, this will require the generation of brain specific conditional knockout models as adult ATF4 germline knockout mice exhibit peripheral phenotypes including decreased bone mass and microphthalmia (defect in eye lens formation that renders them essentially blind) and thus are unsuitable for motor or cognitive behavioural testing (Hettmann, Barton, & Leiden, 2000; Masuoka & Townes, 2002). Interestingly, ATF4 has been implicated in other neurodegenerative conditions but the mechanisms by which it regulates neuronal loss have yet to be fully understood. Specifically, in Alzheimer’s disease, elevated levels of ATF4 has been found in *post-mortem* patient brains and in mice amyloid-beta causes ATF4 production within axons leading to neurodegeneration (Baleriola et al., 2014). Furthermore, in a mouse model of Amyotrophic Lateral Sclerosis (ALS), ATF4-null transgenic ALS animals exhibited delayed onset of disease and displayed a prolonged lifespan compared ATF4-wild type/ALS animals (Matus, Lopez, Valenzuela, Nassif, & Hetz, 2013).

In this study, we demonstrated that ATF4 is required for the transcriptional activation of known pro-apoptotic factors CHOP, Trib3, and the Bcl-2 family member PUMA. Interestingly, all three of these factors have previously been reported to contribute to dopaminergic neuronal death in PD models (Aimé et al., 2015; Bernstein et al., 2011; Bernstein & O’Malley, 2013; Silva et al., 2005). However, the mechanism by which these targets lead to neuronal loss is not completely understood. CHOP and Trib3 have previously been reported to negatively regulate the serine/threonine survival kinase Akt, leading to dephosphorylation and nuclear translocation of different FoxO transcription factors (Du, Herzig, Kulkarni, & Montminy, 2003; Ghosh, Klocke, Ballestas, & Roth, 2012). In an amyloid-beta toxicity study, Saleem and Biswas (2017) described an increase in Trib3 levels in both *in vivo* and *in vitro* experiments which leads to downregulation of Akt and initiation of FoxO1 translocation. We have previously demonstrated that trophic factor deprivation downregulates Akt and promotes FoxO3a mediated transcriptional activation of PUMA suggesting that these apoptotic factors may be involved in a common apoptotic pathway (Ambacher et al., 2012). Indeed, we have previously shown PUMA to be a key regulator of oxidative stress and ER-stress induced neuronal apoptosis (Steckley et al., 2007; Galehdar et al 2010).

The ISR is activated in many neurodegenerative disorders and several studies have investigated the therapeutic potential of targeting eIF2α kinases in mouse models (Hugon, Mouton-Liger, Dumurgier, & Paquet, 2017; Scheper & Hoozemans, 2013). Related to PD it has recently been reported that PERK signalling is elevated in PD brains and in the SNpc of rodents treated with 6-OHDA (Mercado et al., 2018b). Furthermore, they demonstrated that administration of the PERK inhibitor GSK2606414 protected nigrostriatal neurons against 6-OHDA induced toxicity and improved motor function. In this study we found that the PKR inhibitor C16 reduced ATF4 induction and ATF4 mediated transcriptional induction of the pro-apoptotic targets Chop and Trib3. Furthermore, we found that C16 markedly reduced MPP+ and 6-OHDA induced dopaminergic neuron death in mesencephalic cultures. Interestingly, we found that C16 also attenuated ATF4 activation by oxidative stressors and the classic ER stressor thapsigargin. ER stress is generally thought to activate ATF4 via the PERK pathway suggesting that C16 may be able to inhibit PERK in addition to PKR. However, we can also not rule out the possibility that C16 may be affecting ATF4 activation via an off-target mechanism. Indeed, it has previously been reported that C16 can also exert neuroprotection via PKR-independent mechanisms possibly involving inhibition of CDK5 (Chen, Wang, & D’Mello, 2008). Considering this, further studies should aim to better understand the mechanism by which C16 exerts its neuroprotective effects and how this compound may contribute to a reduction in ATF4-induced neuronal cell death.

